# Apoptotic contraction drives target cell release by cytotoxic T cells

**DOI:** 10.1101/2022.11.02.514865

**Authors:** Elisa E. Sanchez, Maria Tello-Lafoz, Aixuan J. Guo, Miguel de Jesus, Benjamin Y. Winer, Sadna Budhu, Eric Chan, Eric Rosiek, Taisuke Kondo, Justyn DuSold, Naomi Taylor, Gregoire Altan-Bonnet, Michael F. Olson, Morgan Huse

## Abstract

Cytotoxic T lymphocytes (CTLs) use immune synapses to destroy infected or transformed target cells. Although the mechanisms governing synapse assembly have been studied extensively, little is known about how this interface dissociates, which is a critical step that both frees the CTL to search for additional prey and enables the phagocytosis of target corpses. Here, we applied time-lapse imaging to explore the basis for synapse dissolution and found that it occurred concomitantly with the cytoskeletal contraction of apoptotic targets. Genetic and pharmacological disruption of apoptotic contraction indicated that it was necessary for CTL dissociation. Furthermore, acute stimulation of contractile forces triggered the release of live targets, demonstrating that contraction is sufficient to drive the response. Finally, mechanically amplifying apoptotic contractility promoted faster CTL detachment and serial killing. Collectively, these results establish a biophysical basis for synapse dissolution and highlight the importance of mechanosensory feedback in the regulation of cell-cell interactions.

## Introduction

Cytotoxic T lymphocytes (CTLs) fight intracellular pathogens and cancer by destroying infected or transformed target cells^1^. Killing responses are typically triggered by the recognition of cognate peptide major-histocompatibility complex (pMHC) on the target cell by T cell antigen receptors (TCRs) on the CTL. This leads to the formation of a specialized cytolytic immune synapse (IS) between the two cells^2, 3^, into which the CTL secretes a toxic mixture of the hydrophobic protein perforin and several granzyme proteases. Perforin forms oligomeric pores in the target cell plasma membrane that enable granzymes to access the cytoplasm^4^, where they induce apoptosis by cleaving caspase-3 and several other substrates^5^. In recent years, CTL-mediated killing has emerged as a critical component of cancer immunotherapy, both in the context of checkpoint blockade^6^, which exploits endogenous TCR specificities, and chimeric antigen receptor (CAR) therapy^7^, which employs a synthetic receptor for a known tumor antigen. As such, there has been a great deal of interest in the mechanisms controlling the formation and maintenance of the cytotoxic IS.

CTLs are highly efficient serial killers that are each capable of destroying ~20 target cells per day in vivo^8, 9^. Serial killing demands not only the precise and expeditious formation of cytolytic synapses but also their timely dissolution so that CTLs can search for additional prey. Effective release of dead or dying target cells is also essential for the clearance of apoptotic debris by professional phagocytes, which minimizes the exposure of surrounding tissue to potentially inflammatory intracellular components^10^. Despite the obvious importance of IS termination for CTL function, the mechanisms that control it are poorly understood. It has been suggested that the release of perforin and granzyme (also called degranulation) is itself the trigger for dissociation^11^. This model, however, is inconsistent with data showing that granzyme and perforin deficient CTLs, which partially or completely lack killing capacity, remain attached to target cells despite degranulating^12^. Indeed, the importance of cytotoxic potential for CTL detachment implies that the process has a CTL-extrinsic trigger, namely damage or death of the target.

How might the IS “sense” damage? The most obvious answer is via recognition of biochemical features characteristic of cellular distress. Apoptosis, for instance, exposes the intracellular lipid phosphatidylserine^13^, which could, in principle, promote IS dissolution by engaging an inhibitory receptor. At present, however, there is no evidence for a mechanism of this kind. Alternatively, dying cells might decrease their expression of activating ligands, becoming less stimulatory to CTLs and thereby encouraging detachment. In support of this model, lymphocytic target cells have been shown to downregulate multiple activating ligands during natural killer (NK) cell-induced apoptosis^14^. It remains unclear, however, if the timing of ligand downregulation matches that of target cell release, and perturbation experiments linking the former process to the latter have not been performed.

Apoptotic cells undergo stereotyped architectural changes, as well, beginning with the rapid contraction of the F-actin cytoskeleton, followed by the formation of blebs and apoptotic bodies^15^. This morphological response represents an intriguing candidate for the induction of target cell release because cytotoxic lymphocytes are both physically dynamic and mechanosensitive^16^. Many critical activating receptors, including the TCR and the α_L_β_2_ integrin LFA-1, require applied force to bind their ligands with high affinity and to transduce downstream signals^17, 18, 19, 20^. This requirement is thought to place physical demands on the target surface, which must be rigid enough to counterbalance the mechanical load placed on receptor-ligand complexes^16^. Consistent with this idea, it has been shown that stiff stimulatory substrates or target cells induce stronger T cell activation than their softer counterparts^21, 22, 23^. Similarly, reducing the lateral mobility of activating ligands within the target membrane increases their stimulatory capacity^24^. Accordingly, it is conceivable that an acute change in the mechanical state of the target cell, independent of it chemical composition, would be sufficient to induce IS dissolution.

In the present study, we employed time-lapse videomicroscopy to examine how CTLs release their targets. Our results reveal that cytoskeletal contraction of the target cell is both necessary and sufficient for IS dissolution, implying that CTLs sense and respond to the morphomechanical indices of programmed cell death.

## Results

### CTL detachment occurs concomitantly with apoptotic contraction

To investigate the determinants of target cell release, we imaged wild type and perforin knockout (*Prf1^-/-^*) OT-1 CTLs together with B16F10 melanoma cells loaded with ovalbumin257-264 (OVA), which is recognized by the OT-1 TCR in the context of the mouse class I MHC H-2K^b^. Propidium iodide (PI), a DNA intercalator, was included to detect large scale disruption of the plasma membrane, a late stage marker of programmed cell death (Fig. 1a). Wild type CTLs displayed robust killing activity in these cocultures (Fig. 1a-b, supplementary movie 1), forming synapses with target cells that were both lethal (killing probability 88%) and transient (mean duration 75.7 ± 8.3 min). By contrast, *Prf1^-/-^* CTLs, which cannot kill B16F10 targets, formed hyperstable synapses, the majority of which persisted for more than 5 hours (Fig. 1c-d, supplementary movie 2). This result, which was in line with prior work^12^, strongly implicated target cell death in the CTL dissociation response. Consistent with this interpretation, we found that the small fraction of wild type CTLs that failed to kill resembled *Prf1^-/-^* CTLs in that they remained attached to their targets for extended periods of time (Fig. 1b).

**Figure 1.**
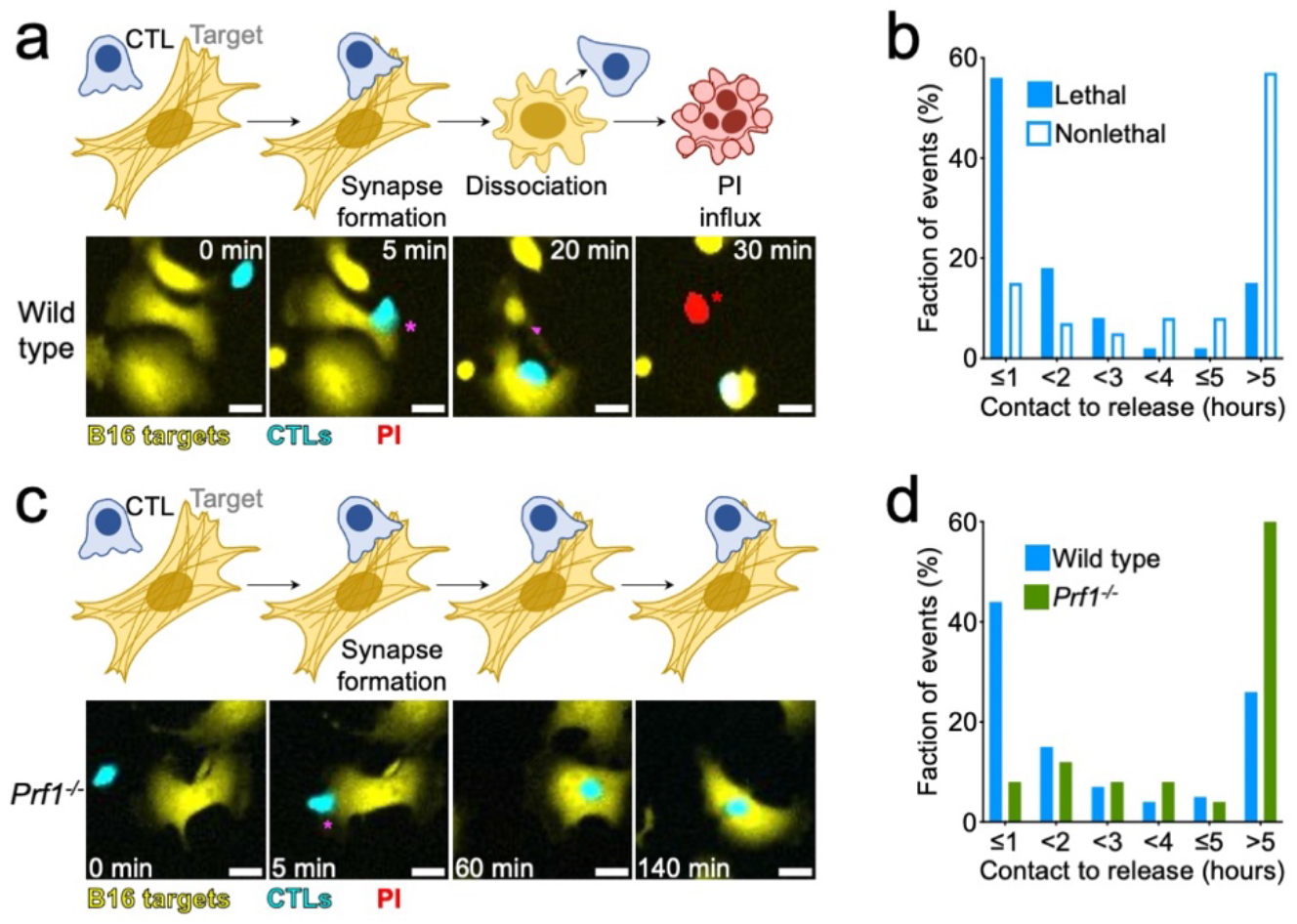
CTL dissociation is triggered by target cell death. CTV-labeled *Prf1^+/+^* and *Prf1^-/-^* OT-1 CTLs were imaged together with OVA-loaded, YFP^+^ B16F10 cells in the presence of PI. (**a** and **c**) Representative time-lapse montages of *Prf1^+/+^* (**a**) and *Prf1^-/-^* (**c**) interactions are shown below, with schematic diagrams above. Magenta asterisks and magenta arrowheads denote IS formation and dissociation, respectively, while red asterisks indicate PI influx. Scale bars = 20 μm. (**b** and **d**) Histograms of IS duration comparing either lethal and nonlethal *Prf1^+/+^* contacts (**b**) or *Prf1^+/+^* and *Prf1^-/-^* contacts (**d**). N = 237 *Prf1^+/+^* conjugates and 191 *Prf1^-/-^* conjugates.

To explore the relationship between target cell death and IS dissolution more closely, we determined the temporal offset between CTL detachment and various steps in the apoptotic cascade. To enable these measurements, we imaged CTL-target cell cocultures in the presence of both PI and CellEvent Caspase-3/7 Green, a probe for global effector caspase activation (Fig. 2a). In B16F10 cells, CTL-mediated killing was characterized by a surge in cytoplasmic CellEvent Caspase-3/7 fluorescence, followed by nuclear PI influx 57.1 ± 3.7 minutes later (Fig. 2a, supplementary movie 3). Before either of these events, however, dying B16F10 cells underwent dramatic contraction of the cell body, transforming from an extended, stellate configuration into a compact spheroid with multiple associated blebs. This contraction occurred rapidly (duration of 15.6 ± 1.4 min), and it represented the first visible sign of target cell distress in the vast majority of cytotoxic conjugates. Based on its timing relative to the other apoptotic indices, we reasoned that this architectural change corresponded to largescale actomyosin-based contraction, an established early event in the apoptotic cascade^15^. Interestingly, contraction coincided with or slightly preceded CTL detachment in a majority of killing events, making it a better predictor of IS dissolution than either CellEvent Caspase-3/7 fluorescence or PI influx, both of which occurred well after detachment in most cases (Fig. 2b). To examine whether this correlation also applied in three dimensional (3D) culture, we imaged mixtures of OT-1 CTLs and antigen-loaded B16F10 targets that had been embedded in collagen-fibrin gels (Supplementary figure 1a). As target cell spreading was less obvious under these conditions, we could not pinpoint the moment of apoptotic contraction for all killing events. Nevertheless, in conjugates where contraction could be scored, it was a better predictor of dissociation than was PI influx (Supplementary figure 1b-c, supplementary movie 4). Hence, target cell contraction is closely associated with CTL release in both 2D and 3D environments.

**Figure 2.**
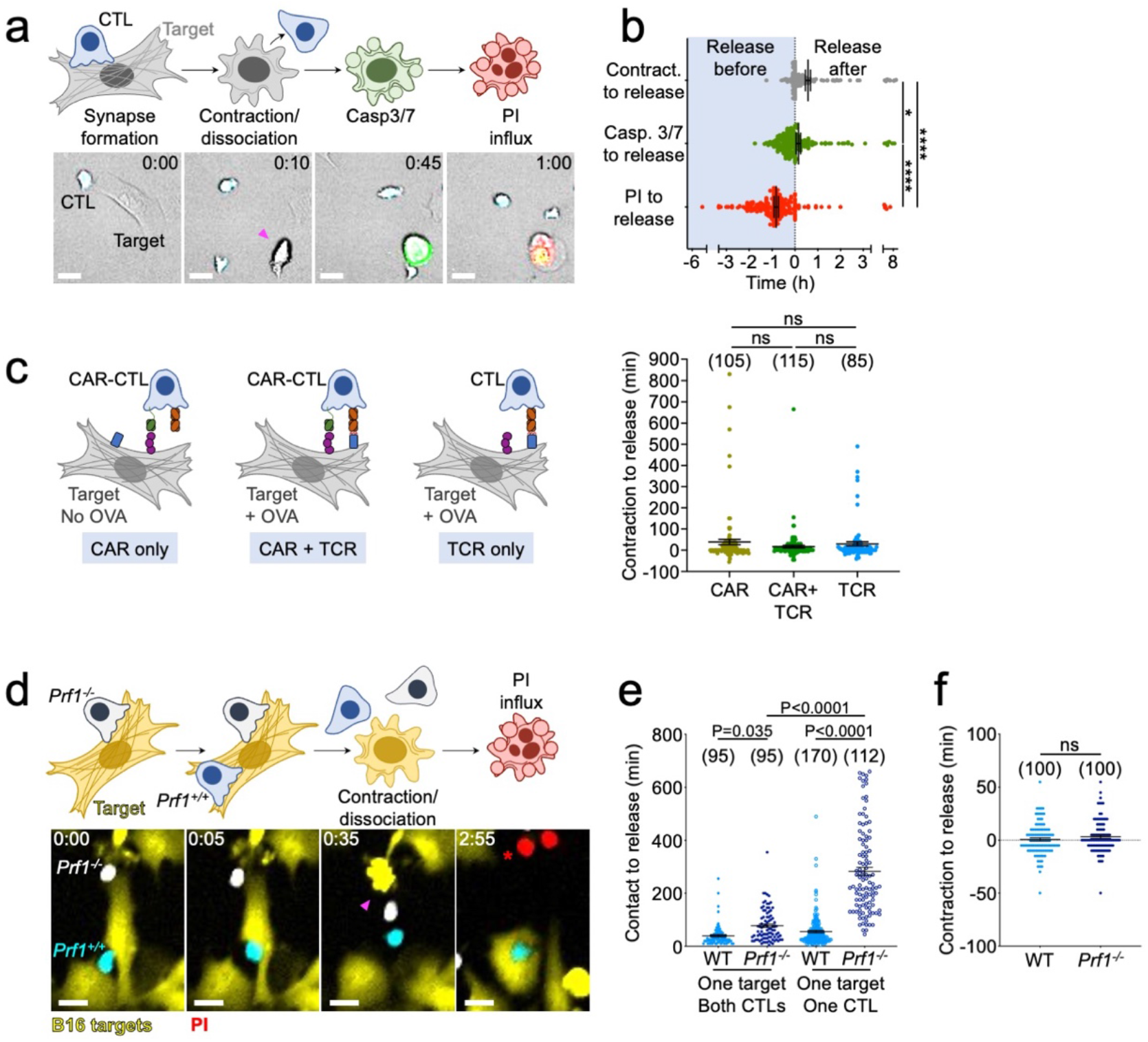
CTL dissociation is temporally correlated with apoptotic contraction. (**a-b**) CTV-labeled OT-1 CTLs were imaged together with OVA-loaded B16F10 cells in the presence of CellEvent Caspase-3/7 and PI. (**a**) A time-lapse montage of a representative contact is shown below, with schematic diagrams above. Dissociation is indicated by the magenta arrowhead. Time in H:MM is shown in the top right corner of each image. (**b**) Quantification of offset time between contraction (Contract.), CellEvent Caspase 3/7 flux (Casp. 3/7), or PI flux and target cell release. Data points to the left of the origin denote detachment before the index of interest, whereas data points to the right of the origin denote detachment after. N = 195. (**c**) CTV-labeled OT-1 CTLs were transduced with either CD19-CAR or empty vector. They were then CTV-labeled and imaged in the presence of CellEvent Caspase-3/7 and PI together with CD19 expressing B16F10 target cells that had been either pulsed with OVA or left unpulsed. Left, a schematic diagram showing CAR and TCR engagement in each of the experimental groups. Right, quantification of the time delay between the onset of contraction and target cell release. (**d-f**) CTV-labeled *Prf1^+/+^* OT-1 CTLs and CellVue Maroon-labeled *Prf1^-/-^* OT-1 CTLs were imaged together with OVA-loaded, YFP^+^ B16F10 in the presence of PI. (**d**) A time-lapse montage of a representative ternary interaction is shown below, with schematic diagrams above. Dissociation and PI influx are indicated by the magenta arrowhead and red asterisk, respectively. Time in H:MM is shown in the top left corner of each image. (**e**) Quantification of IS duration for the indicated *Prf1^+/+^* (WT) and *Prf1^-/-^* CTLs. (**f**) Quantification of the time delay between the onset of contraction and target cell release, for both *Prf1^+/+^* (WT) and *Prf1^-/-^* CTLs involved in ternary complexes. All scale bars = 20 μm. In **c**, **e**, and **f**, sample size is shown in parentheses at the top of each column, and error bars denote SEM. All P-values were calculated by one-way ANOVA with Tukey correction. *, ****, and ns denote P ≤ 0.05, P ≤ 0.0001, and P > 0.05, respectively.

To assess the generality of these observations, we performed imaging studies using two other H-2K^b+^ targets, MB49 bladder cancer cells and C57BL/6 cardiac endothelial cells. CTL detachment was best predicted by apoptotic contraction in each case (supplementary fig. 2), indicating that this correlation is not unique to the B16F10 system. We also examined CAR dependent killing by imaging CD19^+^ B16F10 cells together with OT-1 CTLs expressing a second generation CD19 specific CAR (Fig. 2c). Imaging runs were performed in both the presence and absence of OVA peptide and also using mock-transduced CTLs, enabling us to compare cytotoxic interactions induced by CAR engagement with those induced by the TCR or by the simultaneous engagement of both receptors. CTL detachment was tightly correlated with target cell contraction in all cases (Fig. 2c), indicating that this dissociation mechanism applies to most, if not all, types of cytotoxic synapses. Consistent with this interpretation, we found that murine NK cells also release apoptotic B16F10 cells during the contraction phase (supplementary fig. 3).

We next investigated whether the close association between contraction and detachment was intrinsic to the CTL responsible for killing. Antigen-loaded B16F10 cells were imaged together with both *Prf1^+/+^* and *Prf1^-/-^* CTLs, and ternary interactions containing one target, one *Prf1^+/+^* CTL, and one *Prf1^-/-^* CTL were compared to conjugates containing only one CTL and one target (Fig. 2d). The dynamics of one-to-one conjugates recapitulated the results we obtained from cocultures containing only one type of CTL; *Prf1^-/-^* synapses were long-lived while *Prf1^+/+^* contacts were transient, dissociating concomitantly with apoptotic contraction (Fig. 2e-f). By comparison, in ternary complexes, target cell apoptosis, presumably driven by the *Prf1^+/+^* CTL, was tightly correlated with the detachment of both the *Prf1^+/+^* and the *Prf1^-/-^* CTL (Fig. 2d, supplementary movie 5). This had the effect of dramatically shortening *Prf1^-/-^* CTL contact times in ternary complexes containing *Prf1^+/+^* CTLs (Fig. 2e). Indeed, within ternary complexes, the time intervals between apoptotic contraction and the dissociation of *Prf1^-/-^* CTLs were statistically indistinguishable from the corresponding *Prf1^+/+^* CTL intervals (Fig. 2f). These results indicate that apoptosis drives the dissolution of both cytolytic and non-cytolytic synapses and further implicate contraction as the trigger for the dissociation response.

### Cytoskeletal contraction is necessary for CTL dissociation

Canonical apoptotic contraction is driven by actomyosin-based contractility^15, 25, 26^. To assess the role of the target cell cytoskeleton in the CTL-induced response, we used confocal microscopy to image cocultures of CTLs and antigen-loaded MB49 cells expressing Tractin-RFP, a probe for filamentous-actin (F-actin), and Myl12b-GFP, a probe for nonmuscle myosin II. Concomitant with the onset of contraction, we observed strong accumulation of both F-actin and myosin in cortical cables oriented parallel to the primary axis of morphologic change (Fig. 3a, supplementary movie 6). The fluorescence intensity of these cables increased as the cell contracted (Fig. 3b, supplementary fig. 4a), consistent with the overlap and bundling of F-actin fibers. Interference reflection microscopy of individual killing events revealed that contracting MB49 cells detached completely from the substrate (Supplementary fig. 4b), implying complete disassembly of underlying focal adhesions.

**Figure 3.**
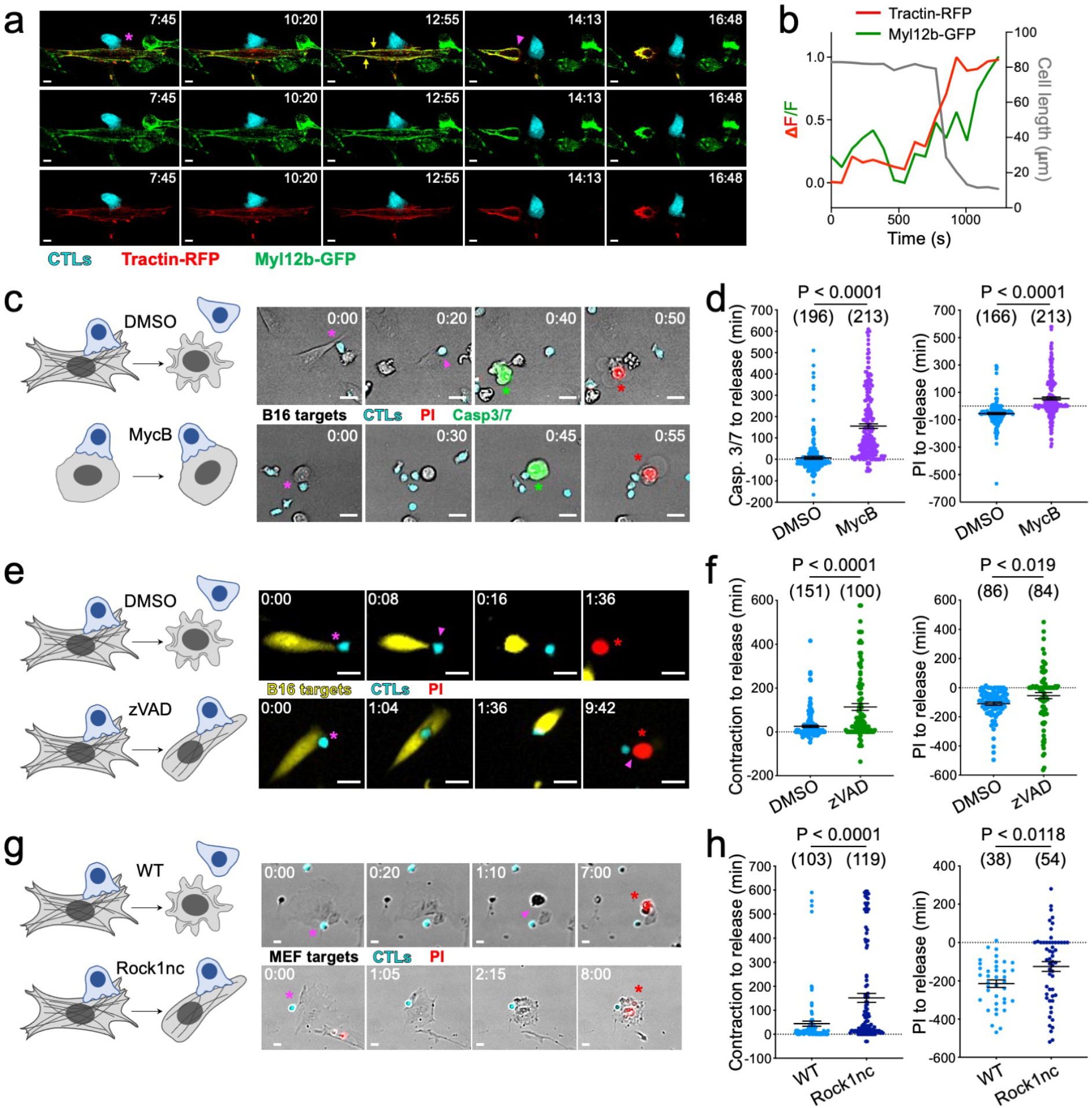
Cytoskeletal contraction is necessary for CTL dissociation. (**a-b**) CTV-labeled OT-1 CTLs were imaged together with OVA-loaded MB49 cells expressing both Myl12b-GFP and Tractin-RFP. (**a**) Time-lapse montages showing a representative dissociation event. All three channels are shown in the top row, while RFP and GFP channels have been removed from the second and third rows, respectively. IS formation is indicated by the magenta asterisk, and dissociation by the magenta arrowhead. Yellow arrows highlight the accumulation of F-actin and myosin in longitudinal cables just prior to contraction. Time in MM:SS is shown in the top right corner of each image. Scale bars = 5 μm. Representative of 6 time-lapse videos. (**b**) Mean fluorescence intensity of Myl12b-GFP and Tractin-RFP in the target cell shown in **a**, graphed over time together with the length of the cell along its major axis. (**c-d**) OVA-loaded B16F10 cells pretreated with MycB or vehicle control (DMSO) were imaged together with CTV-labeled OT-1 CTLs in the presence of CellEvent Caspase-3/7 and PI. (**c**) Time-lapse montages of representative contacts are shown to the right, with schematic diagrams to the left. Conjugate formation is indicated by magenta asterisks, dissociation by the magenta arrowhead, CellEvent Caspase-3/7 flux by green asterisks, and PI influx by red asterisks. Time in H:MM is shown in the top right corner of each image. (**d**) Quantification of the time delays between CellEvent Caspase-3/7 flux (Casp. 3/7) and target cell release (left), and between PI influx and target cell release (right). (**e-f**) CTV-labeled OT-1 CTLs were imaged together with OVA-loaded, YFP^+^ B16F10 cells in the presence of PI and either 100 μm zVAD or vehicle control (DMSO), as indicated. (**e**) Timelapse montages of representative contacts are shown to the right, with schematic diagrams to the left. Conjugate formation is indicated by magenta asterisks, dissociation by the magenta arrowhead, and PI influx by red asterisks. Time in H:MM is shown in the top left corner of each image. (**f**) Quantification of the time delays between the onset of contraction and target cell release (left), and between PI influx and target cell release (right). (**g-h**) CTV-labeled OT-1 CTLs were imaged together with OVA-loaded wild type (WT) or Rock1nc MEFs in the presence of PI. (**g**) Time-lapse montages of representative contacts are shown to the right, with schematic diagrams to the left. Conjugate formation is indicated by magenta asterisks, dissociation by the magenta arrowhead, and PI influx by red asterisks. Time in H:MM is shown in the top left corner of each image. (**h**) Quantification of the time delays between the onset of contraction and target cell release (left), and between PI influx and target cell release (right). Scale bars in **c**, **e**, and **g** = 20 μm. In **d**, **f**, and **h**, sample size is shown in parentheses at the top of each column, and error bars denote SEM. All P-values were calculated by unpaired, two-tailed Student’s t-test.

To evaluate the importance of this cytoskeletal remodeling response for IS dissolution, we sought to decouple it from CTL-mediated killing. The sponge toxin Mycalolide B (MycB) induces near complete actin depolymerization at micromolar concentrations. Importantly, cells treated with MycB remain depleted of F-actin for hours after washout, making it possible to coculture them with CTLs without affecting the CTL cytoskeleton. MycB treatment dramatically altered the architecture of B16F10 cells, transforming them into collapsed, spherical blobs. Importantly, CTLs still formed functional cytotoxic synapses with these targets, inducing robust CellEvent Caspase-3/7 fluorescence and PI influx (Fig. 3c, supplementary movies 7 and 8). Cell death, however, was not accompanied by cytoskeletal contraction, and in the absence of contraction, CTL detachment was markedly delayed. Indeed, most CTLs remained attached to their targets well after global cytoplasmic caspase activation and PI influx (Fig. 3d). To further explore the role of target cell Factin, we transfected B16F10 cells with *Salmonella enterica* SpvB (also called DeAct), an ADP ribosyltransferase that drives dose dependent F-actin severing^27^. Similar to MycB treatment, overexpression of DeAct induced “preemptive” target cell collapse. This prevented normal apoptotic contraction and was associated with an obvious lag in CTL detachment (Supplementary fig. 5a-b, supplementary movies 9 and 10). DeAct expression levels, which we evaluated using a linked Cherry marker, were positively correlated with the duration of contacts (Supplementary fig. 5c), indicating that the degree of cytoskeletal dysfunction dictated the degree of dissociation defect. We also documented an inverse relationship between the starting area of the target cell (a measure of cell spreading) and IS lifetime (Supplementary fig. 5d), suggesting that the capacity of target cells to spread, and therefore to change shape during killing, determined their capacity to induce CTL detachment.

Apoptotic contraction depends in part on caspase-mediated cleavage of Rock1^15, 25^, a serine threonine kinase that induces the activation of myosin II. Caspase processing removes the autoinhibitory C-terminal domain of Rock1, generating a constitutively active protein that drives dysregulated contractility. To evaluate the importance of this pathway for IS dissolution, we first imaged CTL-target cell conjugates in the presence of zVAD-FMK (zVAD), a broad specificity caspase inhibitor. zVAD slowed the apoptotic contraction of target cells (supplementary fig. 5e), but did not prevent their killing (Fig. 3e, supplementary movies 11 and 12), in agreement with prior work showing that blocking caspase activity converts apoptosis into alternative forms of cell death^28, 29, 30^. IS lifetime increased markedly under these conditions (Fig. 3f), corroborating the link between contraction and CTL detachment. We observed the same pattern of results in 3-D collagen-fibrin gels (supplementary fig. 1c), indicating that our results were not tied to specific culture conditions. To interrogate the relevance of Rock1 cleavage more directly, we employed a previously described Rock1 point mutant lacking the caspase recognition motif in its C-terminal tail^31^. Mouse embryonic fibroblasts (MEFs) containing this non-caspase-cleavable form of Rock1 (Rock1nc) knocked into the *Rock1* locus^31^ were OVA-loaded and imaged together with OT-1 CTLs. Rock1nc MEFs contracted more slowly during killing interactions than did wild type MEFs derived from the same genetic background (Fig. 3g, supplementary fig. 5f, supplementary movies 13 and 14). This morphological phenotype was associated with a significant delay in CTL dissociation (Fig. 3h), further supporting the idea that the rate of detachment depends on the strength of the contractile response.

### Cytoskeletal contraction is sufficient for CTL dissociation

Although the studies described above indicate that target cell contraction is necessary for IS dissolution, they do not address whether it is sufficient. For this purpose, we turned to a conditionally activatable Rock construct containing the Rock2 kinase domain, which targets the same substrates as Rock1, fused to the hormone binding domain of the estrogen receptor (ER)^32^ (Fig. 4a). The Rock2-ER fusion dimerizes in the presence of 4-hydroxytamoxifen (4-OHT, an estrogen antagonist), leading to kinase activation and actomyosin contractility in the absence of cell death. Transient transfection of MB49 cells with GFP-Rock2-ER yielded a range of morphological phenotypes at a single cell level. Many of the high expressors were constitutively rounded up, implying 4-OHT independent Rock signaling. The lower expressors spread normally, however, and a substantial fraction of these cells (81%) exhibited contraction after 4-OHT treatment, with a mean offset time of 69 minutes. MB49 cells expressing a control construct (GFP-ER) did not respond to 4-OHT in this way (10% contraction over 12 h), validating the specificity of our approach.

**Figure 4.**
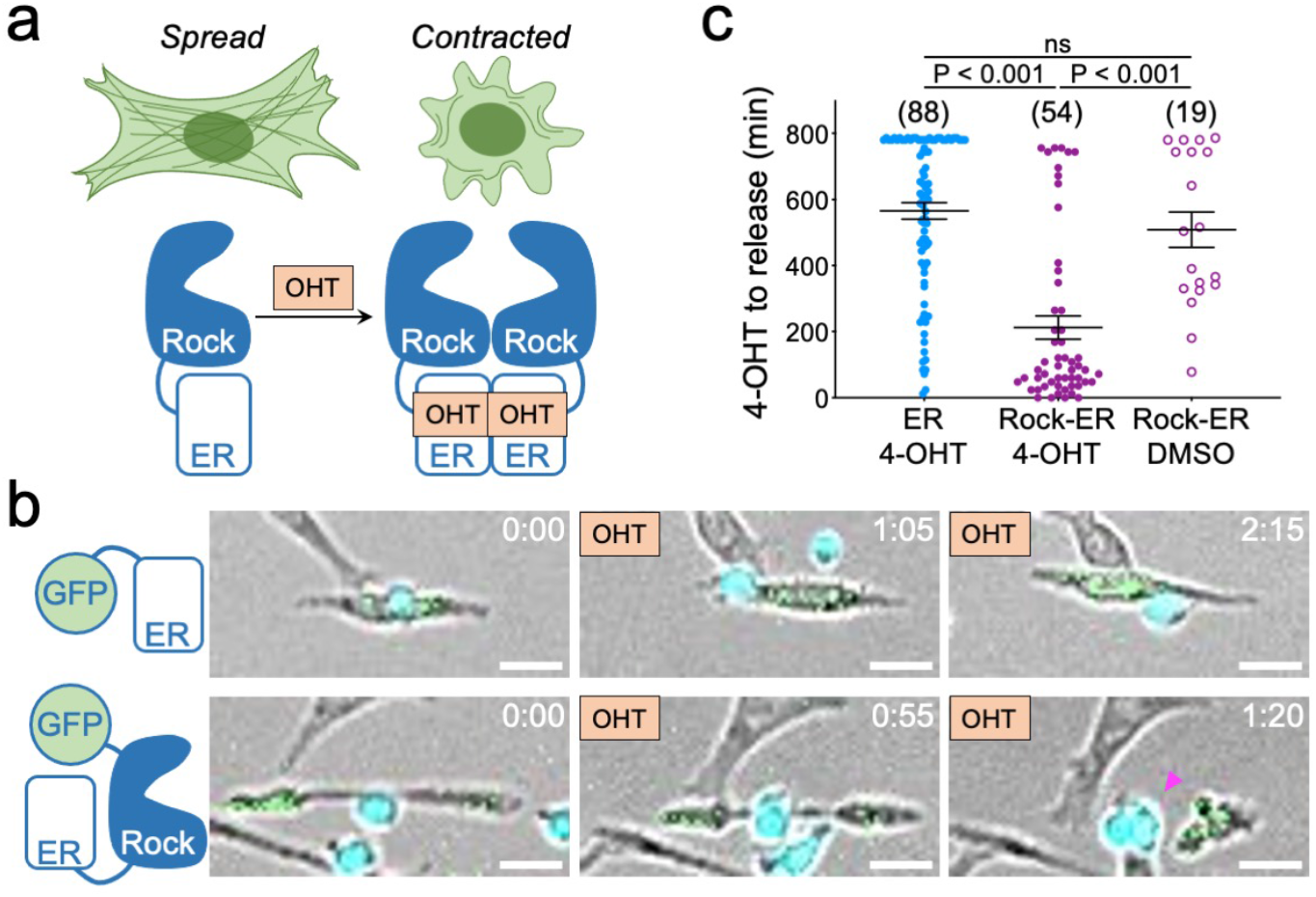
Cytoskeletal contraction is sufficient for CTL dissociation. MB49 cells expressing GFP-Rock2-ER or GFP-ER control were loaded with OVA and imaged together with CTV-labeled *Prf1^-/-^* OT-1 CTLs. (**a**) Treatment with 4-OHT (OHT) induces the dimerization and activation of Rock2-ER, which in turn promotes actomyosin contractility. (**b**) Time-lapse montages showing the effects of 4-OHT on representative GFP-ER (top) and GFP-Rock2-ER (bottom) target cells engaged with CTLs. “OHT” indicates the addition of 4-OHT, and the magenta arrowhead denotes dissociation. Time in H:MM is shown in the top right corner of each image. Scale bars = 20 μm. (**c**) Quantification of the time delay between 4-OHT treatment and target cell release. Sample size is shown in parentheses at the top of each column, and error bars denote SEM. P-values were calculated by one-way ANOVA with Tukey correction. ns = not significant.

With these modified target cells in hand, we next performed imaging studies to determine whether Rock2-ER dependent cytoskeletal remodeling affects IS stability. To minimize the incidence of apoptosis-induced contraction in these experiments, we employed only OT-1 *Prf1^-/-^* CTLs, which cannot kill MB49 cells, and we also treated cocultures with zVAD. 4-OHT was typically added after 2.5 h of a 16 h time-lapse. For analysis, we focused on the CTLs that formed conjugates with GFP^+^ target cells prior to the addition of 4-OHT. In GFP-Rock2-ER cultures, 69% of these CTLs detached within 3 hours of 4-OHT addition (Fig. 4b-c, supplementary movie 15). Interactions between CTLs and GFP-ER targets were much more stable by comparison, with 70% persisting for longer than 8 hours post-treatment (Fig. 4b-c, supplementary movie 16). Closer examination of the GFP-Rock2-ER conjugates revealed that the vast majority of dissociation events occurred in conjugates where target cell contraction actually occurred. Indeed, vehicle treated GFP-Rock2-ER target cells, which did not contract, exhibited IS stability that was statistically indistinguishable from that of 4-OHT treated GFP-ER controls (Fig. 4c). These results strongly suggest that Rock dependent cytoskeletal contraction is, independently of cell death, sufficient to trigger IS dissolution.

### CTLs detect target cell contraction through the IS

Apoptotic contraction could disrupt the IS directly by tearing the target cell away from the CTL. Alternatively, the CTL might sense structural change in the contracting target and then reduce its synaptic avidity to facilitate release (Fig. 5a). To explore these two mechanisms, which are not mutually exclusive, we monitored intracellular calcium (Ca^2+^), an index of T cell activation, during IS formation, target contraction, and release. Consistent with previous studies^33, 34, 35^, we observed that CTL Ca^2+^ levels increased rapidly upon initial contact with a target cell and remained high prior to the onset of apoptosis (Fig. 5b, supplementary movie 17). Ca^2+^ responses tended to decline, however, during the period between contraction and release, before falling even further after CTL detachment (Fig. 5b-c, supplementary movie 17). By graphing target cell area measurements with paired CTL Ca^2+^ data from each IS as individual trajectories, we confirmed that target contraction (visualized as smaller area measurements) was indeed associated with reduced Ca^2+^ signaling (Fig. 5d). To further investigate whether these lower Ca^2+^ levels were caused by contraction itself, or simply reflected a time dependent decline in Ca^2+^ signaling, we compared wild type responses with those of *Prf1^-/-^* CTLs, which do not induce B16F10 contraction or death. *Prf1^-/-^* responses did not decline as soon or as steeply as their wild type counterparts, indicating that a sharp fall-off in Ca^2+^ levels is a characteristic property of bona fide cytolytic interactions. Notably, the separation between the wild type and *Prf1^-/-^* response curves became statistically significant only after the majority of target cells in the wild type synapses had begun to contract (indicated by the red CDF line in Fig. 5c). This observation lends further support to the idea that contraction attenuates activating signals in attached CTLs. We conclude that CTLs do indeed sense target cell contraction, implying that they play an active role in the dissociation response.

**Figure 5.**
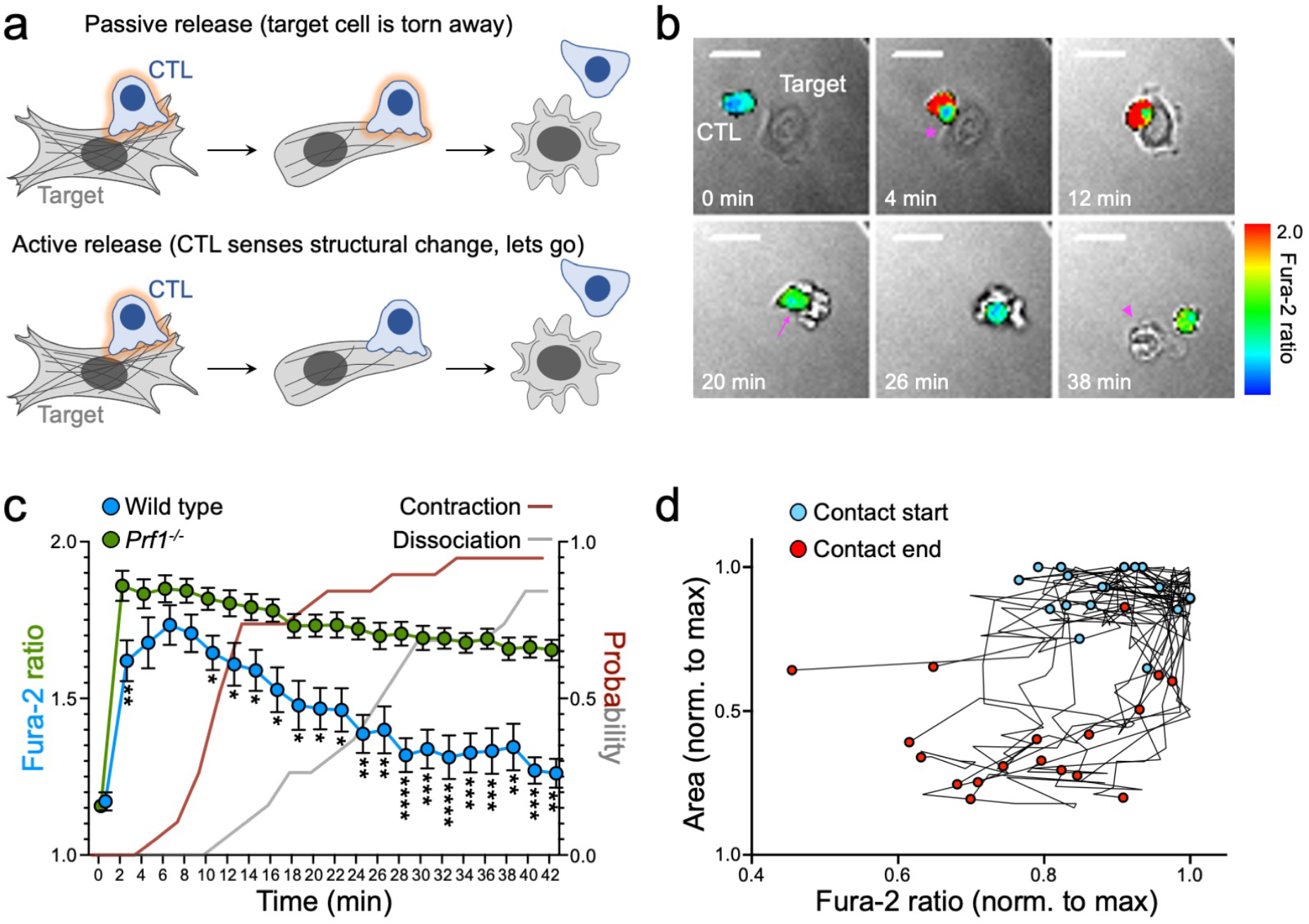
CTLs sense apoptotic contraction through the IS. (**a**) Schematic diagram comparing passive and active mechanisms of contraction-induced CTL release. Activating CTL signaling is indicated by the orange glow around the CTL. (**b-d**) OT-1 CTLs were loaded with Fura-2AM and imaged with OVA-loaded B16F10 cells. (**b**) Time-lapse montage of a representative interaction. Fura-2 ratio is depicted in pseudocolor, with warmer colors indicating higher intracellular Ca^2+^ concentration. Conjugate formation is indicated by the magenta asterisk and dissociation by the magenta arrowhead. Target cell contraction is accompanied by a decrease in intracellular Ca^2+^, denoted by the magenta arrow. Scale bars = 20 μm. (**c**) Mean Ca^2+^ response curves derived from N = 18 wild type responses and N = 43 *Prf1^-/-^* responses. The onset of target cell contraction and CTL dissociation in the wild type conjugates is indicated by the red and gray cumulative distribution functions, respectively. Error bars on Ca^2+^ curves denote SEM. *, **, ***, and **** denote P ≤ 0.05, P ≤ 0.01, P ≤ 0.001, and P ≤ 0.0001, respectively, calculated by unpaired t-test. (**d**) Wild type Ca^2+^ responses from **c** plotted as two dimensional trajectories of Fura-2 ratio against target cell area. Values and the beginning and end of each contact are highlighted in blue and red, respectively.

### Modulating target cell contraction to control CTL detachment and serial killing

Given the importance of efficient IS turnover for serial killing, we speculated that it should be possible to boost CTL function by augmenting the contraction-induced dissociation response. To explore this idea, we employed the myocardin related transcription factors A and B (MRTF-A and MRTF-B), which drive the production of actin, F-actin binding proteins, and other cytoskeletal regulators^36^. Cell lines overexpressing MRTFs contain copious F-actin stress fibers and exhibit increased cortical stiffness relative to unmodified controls^37^. We reasoned that this enhanced cytoskeletal state would effectively amplify the architectural change induced by apoptotic contraction, which would, in turn, drive a stronger dissociation response (Fig. 6a). Consistent with this prediction, we observed a marked acceleration in the rate of CTL detachment from B16F10-MRTF-A and B16F10-MRTF-B cells relative to B16F10 controls (Fig. 6a-b). Importantly, this increase in synaptic turnover was associated with improved serial killing; in cocultures containing B16F10-MRTF-A or B16F10-MRTF-B cells, a substantially larger fraction of CTLs destroyed multiple targets (Fig. 6c).

**Figure 6.**
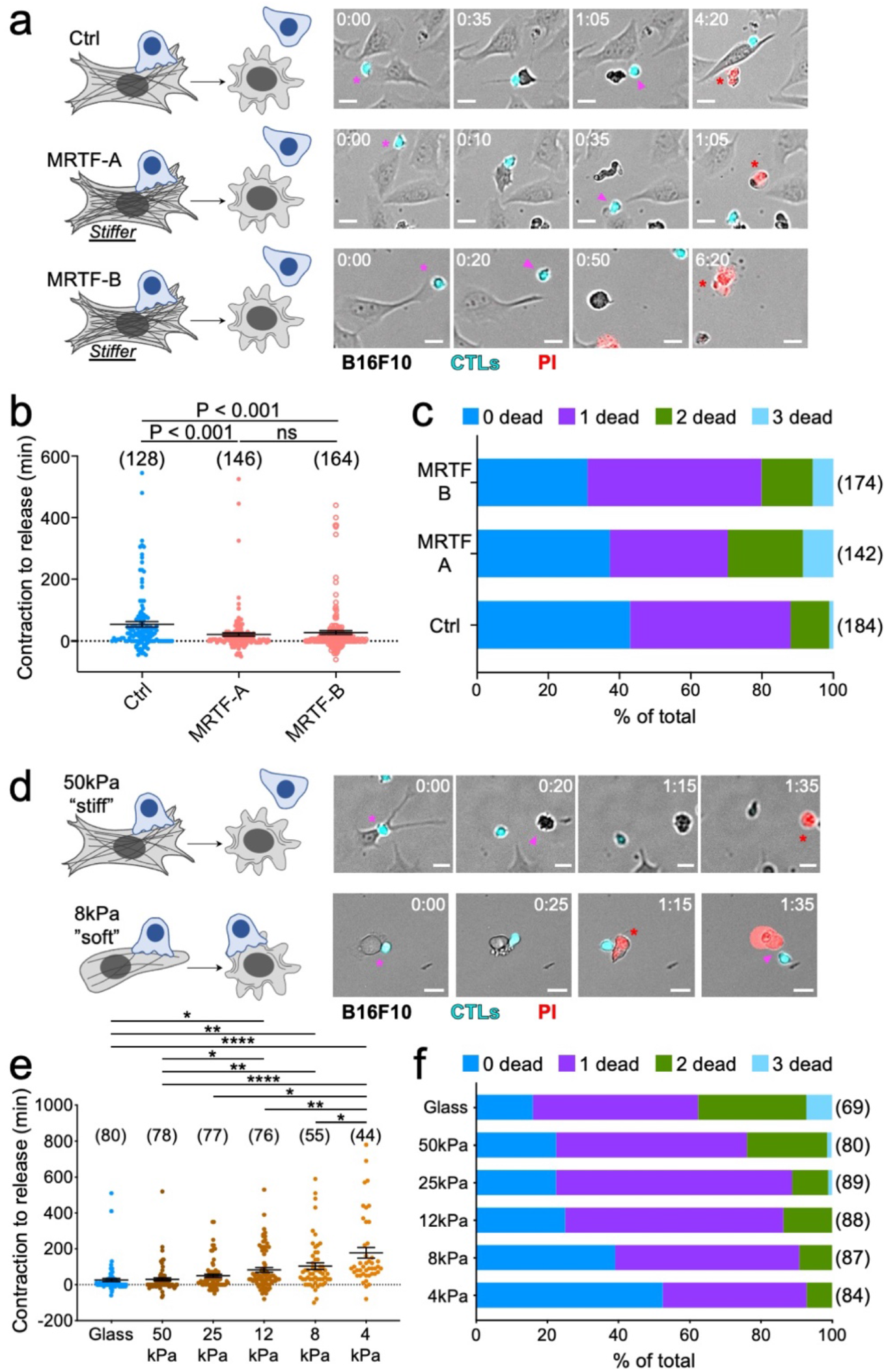
Morphological change during apoptotic contraction controls CTL dissociation and serial killing. (**a-c**) CTV-labeled OT-1 CTLs were imaged in the presence of PI together with OVA-loaded control (Ctrl) B16F10 cells or B16F10 cells overexpressing the indicated MRTF isoforms. (**a**) Time-lapse montages of representative contacts are shown to the right, with schematic diagrams to the left. Conjugate formation is indicated by magenta asterisks, dissociation by magenta arrowheads, and PI influx by red asterisks. Time in H:MM is shown in the top left corner of each image. (**b**) Quantification of the time delay between the onset of contraction and target cell release. (**c**) Cumulative bar graph showing the relative frequency of serial killing in cocultures containing the indicated target cells. (**d-f**) CTV-labeled OT-1 CTLs were imaged in the presence of PI together with OVA-loaded B16F10 cells on hydrogels of varying stiffness or glass. (**d**) Time-lapse montages of representative contacts on 50 kPa and 8 kPa hydrogels are shown to the right, with schematic diagrams to the left. Conjugate formation is indicated by magenta asterisks, dissociation by magenta arrowheads, and PI influx by red asterisks. Time in H:MM is shown in the top right corner of each image. (**e**) Quantification of the time delay between the onset of contraction and target cell release. (**f**) Cumulative bar graph showing the relative frequency of serial killing in cocultures containing the indicated target cells. Scale bars in **a** and **d** = 20 μm. Sample size is shown in parentheses at the top of each column in **b** and **e**, and to the right of each bar in **c** and **f**. Error bars denote SEM. P-values were calculated by one-way ANOVA with Tukey correction. *, **, and **** denote P ≤ 0.05, P < 0.01, and P < 0.0001, respectively.

Adherent cells are modulated continuously by cell-extrinsic biophysical signals. They are particularly responsive to the mechanical properties of the extracellular matrix, which influence the structure and rigidity of the cortical cytoskeleton^38^. Accordingly, it has been shown that cells grown in stiffer environments become stiffer themselves, whereas those in softer environments become more compliant^39, 40^. Consistent with these prior results, we found that B16F10 cells cultured on very stiff fibronectin-coated glass (1000 megaPa Young’s modulus) adopted a spread, stellate morphology containing visible F-actin stress fibers, while the same cells grown on soft hydrogels (e.g. 4 kPa Young’s modulus) were more rounded and exhibited less obvious cytoskeletal structures (supplementary Fig. 6a-b). Stiffer hydrogels (e.g. 50 kPa Young’s modulus) elicited an intermediate phenotype characterized by pronounced extension along one axis. To determine whether these morphological differences affected killing dynamics, B16F10 cells cultured on glass or on hydrogels of varying stiffness were loaded with OVA and then imaged together with OT-1 CTLs. Target cell killing was observed on all substrates, indicating that the fundamental cytotoxic capacity of CTLs was unaffected by the substrate. The quality of apoptotic contraction, however, differed dramatically between conditions. Target cells on glass or stiff hydrogels contracted robustly, whereas cells on soft substrates exhibited a much less obvious response, presumably because their resting state was already somewhat collapsed (Fig. 6d). Muted contraction on soft substrates was accompanied by a marked delay in IS dissolution, with CTLs often remaining attached to their targets well after PI influx (Fig. 6d-e; supplementary fig. 6c). CTLs on soft hydrogels also exhibited less serial killing than their counterparts in stiffer cultures (Fig. 6f). Taken together with the MRTF experiments above, these results confirm the importance of apoptotic contraction for IS dissolution, and they demonstrate that timely target cell release is critical for effective serial killing.

## Discussion

Taken together, our results support a model in which apoptotic contraction, mediated by actomyosin forces downstream of Rock signaling, induces the release of dying target cells by CTLs (Supplementary fig. 7). This model, which is consistent with prior work implicating target cell death in IS dissolution^12^, implies that CTLs continuously monitor the health of their prey through the IS. From a design perspective, conditioning dissociation on target cell death rather than a CTL-intrinsic event, such as degranulation, is preferable because it enables the CTL to adapt in real time to hardier target cells that require multiple degranulation cycles for destruction. Furthermore, basing the death sensing mechanism on an architectural change common to all apoptotic cells, rather than a specific chemical event, ensures that target release is not tied to the formation or suppression of specific receptor-ligand interactions. As a result, the entire process becomes more generalizable. Indeed, contraction-induced dissociation appears to govern NK cells and CAR T cells, as well, indicating that it may be a universal feature of cellular cytotoxicity. The close correlation between target contraction and CAR T cell detachment is particularly notable considering that CARs bind much more tightly than TCRs to their cognate ligands. Thus, contraction-induced dissociation is robust enough to overcome the widespread engagement of activating receptors at the IS, even if those receptors bind with nanomolar affinity.

Inhibiting caspase cleavage of Rock1 delayed CTL-induced contraction but did not prevent it entirely, implying the existence of compensatory mechanisms. Many cell types, including the MEFs used for our Rock1nc experiments, also express Rock2, which lacks a consensus caspase recognition motif but can be cleaved and activated by granzyme B^41^. Accordingly, it is tempting to speculate that the residual apoptotic contraction of Rock1nc MEFs during CTL-mediated killing resulted from direct granzyme B targeting of Rock2. This cleavage event could also explain the diminished, but nevertheless detectable B16F10 contraction response we observed in the presence of zVAD. That both caspase and granzyme dependent pathways have evolved to trigger contraction redundantly suggests that this process is critical for the proper function of cytotoxic lymphocytes.

Our Ca^2+^ imaging experiments indicate that target cell contraction attenuates CTL activation prior to detachment. This reduction in signaling likely reflects the mechanosensitive features of activating immunoreceptors, many of which respond both to the mobility of their cognate ligands in the plasma membrane and to the underlying stiffness of the ligand presenting surface^16^. The β2 integrin LFA-1, for example, achieves optimal ligand binding and outside-in signaling only when it is placed under tension^19, 42, 43^. Dendritic cells exploit this property by restricting the lateral diffusion of LFA-1 ligands on their surface, thereby enhancing their capacity to trigger LFA-1 mechanotransduction and T cell activation^24^. One could imagine apoptotic contraction having the opposite effect by rapidly collapsing the array of activating ligands on the target cell, leading in turn to an acute disruption of CTL mechanotransduction.

A role for mechanotransduction in IS dissolution is consistent with the emerging conception of cellular cytotoxicity as a mechanically active process^16^. Cytotoxic lymphocytes use synaptic forces to establish strong receptor-ligand interactions^17, 19, 20^, position degranulation events within the IS^44^, and potentiate perforin pore formation^45^. Reciprocally, the biophysical disposition of target cells has been shown to dictate both their capacity to elicit lymphocyte activation as well as their sensitivity to perforin^23, 24, 37, 45^. It is also becoming clear that mechanical properties such as cell stiffness respond not only to cell-intrinsic pathways but also to cell-extrinsic crosstalk with the microenvironment^38, 39, 40^. When viewed within this conceptual framework, our data demonstrating that substrate rigidity modulates CTL dissociation rate strongly suggests that microenvironmental mechanics can control the efficiency of cytotoxic interactions. It will be interesting to explore this idea further using biophysically diverse in vivo systems.

Although our results clearly implicate biomechanical change as a trigger for CTL dissociation, they do not rule out a more biochemically-oriented mechanism in which cell death drives target release by altering the composition of activating and inhibitory ligands within the IS. Indeed, it has been shown that dying target cells downregulate certain activating ligands and adhesion molecules^14^, which would be expected to weaken interactions with cytotoxic lymphocytes. Whether ligand downregulation on targets occurs before or after IS dissolution has not been investigated, however, and as such a cause and effect relationship remains to be established for this model. Nevertheless, one can easily imagine scenarios in which both biophysical and biochemical mechanisms are involved in target release. We feel that apoptotic contraction is more likely to be the acute trigger because its kinetics match those of CTL detachment so well. In this regime, ligand downregulation would contribute later in the process by preventing the reengagement of dead targets. Contraction could also inhibit reengagement by reducing the accessible surface area of the apoptotic corpse, effectively making it more difficult to find.

Apoptosis is generally presented as an “altruistic cell suicide” designed to minimize deleterious effects on surrounding tissue^46^. Thus, apoptotic cells assiduously degrade their potentially inflammatory nucleic acids, release anti-inflammatory mediators, and upregulate find-me and eat-me signals to promote phagocytic uptake. Apoptotic contraction, for its part, promotes tissue folding during development and is thought to maintain the barrier integrity of epithelial tissues by facilitating corpse extrusion^47, 48, 49^. Our finding that this morphological change also triggers cytotoxic lymphocyte dissociation is entirely consistent with apoptotic altruism. By letting the killer cell know that its work is done, the target uses its final act to boost the overall efficiency of cellular immunity.

## Methods

### Primary cells and retroviral transduction

To generate CTLs, naïve *Prf1^+/+^* or *Prf1^-/-^* CD8^+^ T cells derived from transgenic mice expressing the OT-1 αβTCR were mixed with splenocytes isolated from C57BL/6 mice that had been pulsed with 100 nM OVA in RPMI medium containing 10% FBS. 30 U/mL of IL-2 was added to cells 24 h following T cell activation. Subsequently, cells were split as needed into RPMI + 10% FBS containing 30 U/mL IL-2 and used for functional assays 6-7 days later. Murine CD19-CAR T cells were prepared by retroviral transduction of OT-1 CTLs 48 h after initial stimulation with OVA. Retrovirus was generated by transient transfection of Phoenix E cells with an MSCV-based expression vector containing a second generation CD19 specific CAR with CD28 and CD3ζ signaling motifs. Virus was collected 48 hours after transfection and used to infect OT-1 CTL blasts via spinfection at 1400 × g in the presence of 4 μg/mL polybrene. Murine NK cells were isolated from C57BL/6 splenocytes by negative selection using an NK cell isolation kit (MACS) followed by overnight incubation in 1000 U/ml IL-2.

### Cell lines

B16F10, MB49, and C57BL/6 cardiac endothelial cells were cultured in RPMI medium containing 10% FBS. ROCK1nc knockin MEFs and wild type control MEFs were cultured in DMEM medium containing 10% FBS. MB49 cells stably expressing Myl12b-GFP were prepared by retroviral transduction. Retrovirus was generated by transient transfection of Phoenix E cells with an MSCV-based expression vector encoding Myl12b-GFP and a puromycin resistance cassette. Following spinfection at 1400 × g in the presence of 4 μg/mL polybrene, transduced MB49 cells were selected by culturing in 2.5 μg/mL puromycin, followed by FACS. Tractin-mCherry was introduced into MB49-Myl12b-GFP cells by transient transfection using lipofectamine 2000 (ThermoFisher), following the manufacturer’s recommended protocol. Inducible MRTF-A and MRTF-B overexpression cell lines (B16F10-MRTF cells) were prepared by sequential transduction with rTTA, TGL, and either control or MRTF overexpression vectors, followed by culture in G418 and hygromycin. MRTF expression was induced by treating cells in 500 ng/ml doxycycline hyclate (Sigma) 24-48 hours prior to the experiment. Cell lines expressing Cherry-DeAct or Cherry alone were prepared 48 hours before the experiment by transient transfection of the DeAct or Cherry control plasmids using lipofectamine 2000. Lipofectamine 2000 was also used to prepared MB49 cells expressing GFP-Rock2-ER or GFP-ER.

### Pharmacological reagents

For depolymerization of the F-actin cytoskeleton, B16F10 target cells were pretreated for 1 h with 100 nM MycB and then washed in RPMI + 10% FBS prior to use. Caspase inhibition was achieved by pretreating B16F10 target cells with 100 μM zVAD for 30 minutes.

### Live imaging of 2D cultures

In preparation for widefield imaging of CTL-mediated killing, target cells (B16F10, MB49, MEF, or cardiac endothelial cells) were cultured overnight on fibronectin-coated 8-well chambered coverglass (Lab-tek) and then pulsed with 100 nM of OVA for 2 h. After washing, targets were mixed with CTV-labeled and/or CellVue Maroon-labeled OT-1 CTLs in the presence of 2 μM Cell Event Caspase-3/7 Green and/or 1.5 μM PI. Chambers containing cells of interest were imaged at 5-8 min intervals using a Zeiss Zen epifluorescence microscope at 10× magnification for 11 hours. IRM was performed on the same Zeiss Zen system fitted with a 50/50 Vis beam splitter and a 63×/1.4 NA objective lens. For Rock2-ER studies, 4-OHT was added after 2.5 h of imaging to a final concentration of 100 μM. In certain experiments, targets were cultured, antigen-loaded, and mixed with CTLs in 96-well glass plates containing fibronectin-coated polyacrylamide hydrogels of different stiffnesses (Matrigen). For CAR T cell studies, B16F10 cells stably transduced with CD19 were used as targets. Widefield Ca^2+^ imaging was performed using the same fibronectin-coated chambered coverglass approach. On the day of imaging, CTLs were loaded with 5 μg/ml Fura-2AM and washed following one hour of staining. Fura-2AM-loaded CTLs were then mixed with adhered OVA-loaded B16F10 targets and imaged with 340 nm and 380 nm excitation at 2 minute intervals for 2 hours, using a 20× objective lens (Olympus). Confocal time lapse videos were acquired on a Leica Sp8 machine equipped with a white light excitation laser and a 40× objective lens. Images were taken every 3 minutes for 2 hours.

### 3D collagen culture

Collagen cultures were formed by resuspending B16F10-YFP target cells (previously pulsed for 2 h in 1 μM OVA) and CTV-labeled OT-1 CTLs in a mixture of collagen (1 mg/mL), fibrinogen (1 mg/mL), and thrombin (20 U/mL) in PBS. Gel mixtures were allowed to harden in 8-well chambered coverglass (Lab-tek) for 20 min. Media containing 100 μM zVAD (or DMSO vehicle) and 1.5 μM PI was then added and the samples incubated for an additional 4-6 h. Subsequently, wells were imaged at 7 min intervals using a Zeiss Zen epifluorescence microscope at 10× magnification for 24 h.

### Fixed imaging

B16F10 cells were cultured overnight on fibronectin-coated hydrogels bound to a glass slide with a removable 8 well media chamber (Matrigen Soft Slide). Cells were then fixed with 2% PFA, permeabilized using 0.5% Triton, stained with Phalloidin Alexa Fluor 568, and imaged using a Zeiss Zen epifluorescence microscope fitted with a 20×/0.4 NA objective.

### Image analysis

To analyze killing kinetics, time intervals between IS formation, cytoskeletal contraction, caspase 3/7 flux, and PI flux were derived from videomicroscopy data. Cytoskeletal contraction was defined as a dramatic and irreversible decrease in cell area with complete loss of morphological asymmetry. Caspase 3/7 and PI responses were scored from the onset of detectable fluorescence. To assess serial killing, the number of targets killed per CTL was calculated. Note that all contacts made between one lymphocyte and one target cell were scored, excluding contacts formed within 30 min of the end of the time-lapse. To quantify DeAct and Cherry expression in target cells, the average fluorescence intensity within cell borders was established using ImageJ software. The same procedure was used to calculate the area of each target cell. Ca^2+^ flux responses (Fura-2 ratios) were quantified using SlideBook (Intelligent Imaging Innovations) and then analyzed in Microsoft Excel. To compare wild type and *Prf1^-/-^* Ca^2+^ responses (Fig. 5c), we ran t-test at every time point for the null hypothesis that the 2 sets of cells (wild type and *Prf1^-/-^*) had identical average ratios of Fura-2. We did not assume that the populations had identical variances. P-values were generated using the scipy.stats.ttest_ind python library. Confocal videos of contracting target cells were analyzed in Imaris (Oxford Instruments) by first calculating the mean fluorescence intensities of Myl12b-GFP and Tractin-RFP within the cell envelope (determined by intensity thresholding in the Myl12b-GFP channel) and then referencing those values to target cell length along its major axis (Fig. 3b and supplementary fig. 4a).

## Acknowledgments

We thank C. Jeronimo for technical support; the MSKCC Molecular Cytology Core Facility for assistance with imaging; M. Overholtzer, M. Baylies, and members of the M.H. laboratory for advice. Supported in part by the U.S. National Institutes of Health (R01-AI087644 to M.H. and E.E.S., R21-AI169847 to M.H., and P30-CA008748 to MSKCC), the Canadian Institutes of Health Research (PJT-169125 to M.F.O.), the Canada Research Chairs Program (950-231665 to M.F.O.), the Ludwig Foundation for Cancer Immunotherapy (M.D.J.), the Schmidt Science Fellows Program (B.Y.W.), the Cancer Research Institute (B.Y.W.), and the Ludwig Institute for Cancer Research (M.T.-L.).

## Author contributions statement

E.E.S., M.T.-L., M.D.J., B.Y.W., S.B., and M.H. designed the experiments.

E.E.S., M.T.-L., A.J.G., E.C., E.R., and M.H. collected the data.

E.E.S., A.J.G., G.A.-B., and M.H. analyzed the data.

S.B., T.K., J.D., N.T., G.A.-B., and M.F.O. contributed key reagents and methods.

E.E.S. and M.H. wrote the paper.

---

**Supplementary figure 1.**
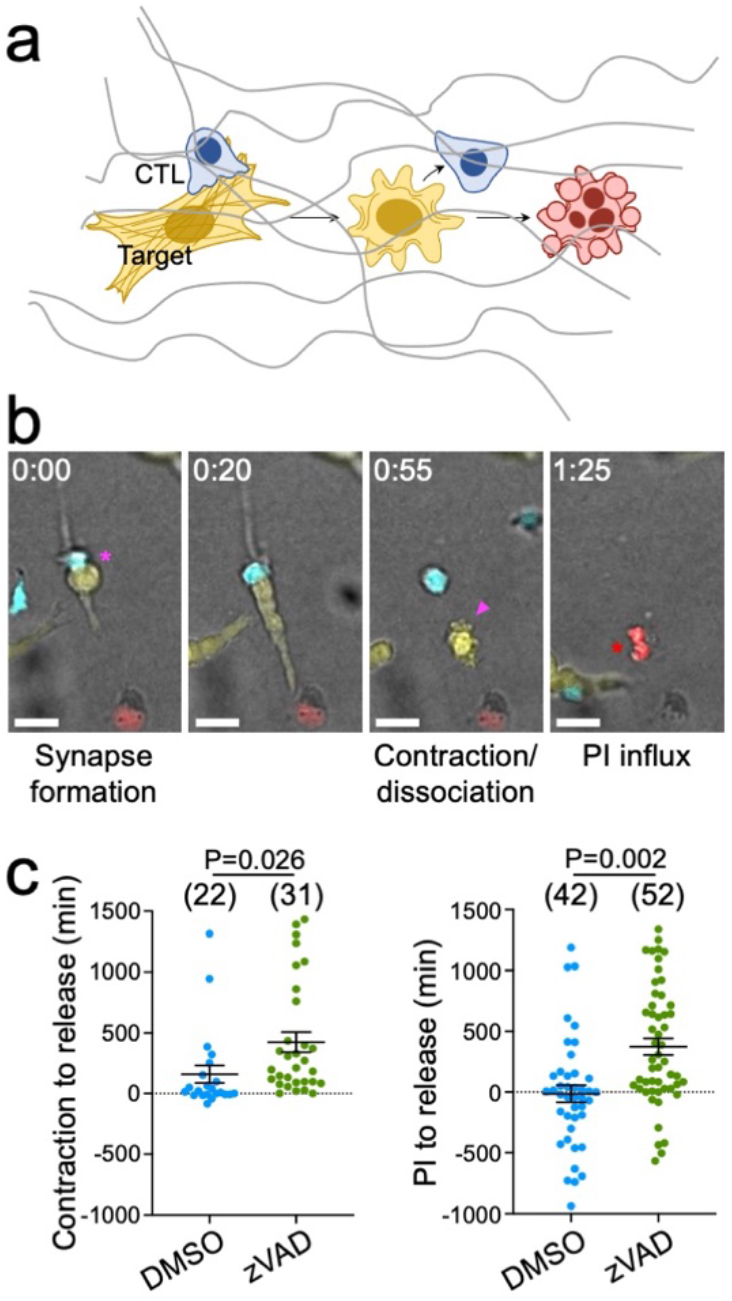
Apoptotic contraction predicts CTL dissociation in 3D culture. Collagen gels containing CTV-labeled OT-1 CTLs and OVA-loaded, YFP^+^ B16F10 cells were imaged in the presence of PI together with either 100 μm zVAD or vehicle control (DMSO), as indicated. (**a**) Diagram schematizing the approach. (**b**) Time-lapse montage of a representative contact, with conjugate formation indicated by the magenta asterisk, dissociation by the magenta arrowhead, and PI influx by the red asterisk. Time in H:MM is shown in the top left corner of each image. Scale bars = 20 μm. (**c**) Quantification of the time delay between the onset of contraction and target cell release (left) and between PI influx and target cell release (right). Sample size is shown in parentheses at the top of each column, and error bars denote SEM. P-values were calculated by unpaired, two-tailed Student’s t test.

**Supplementary figure 2.**
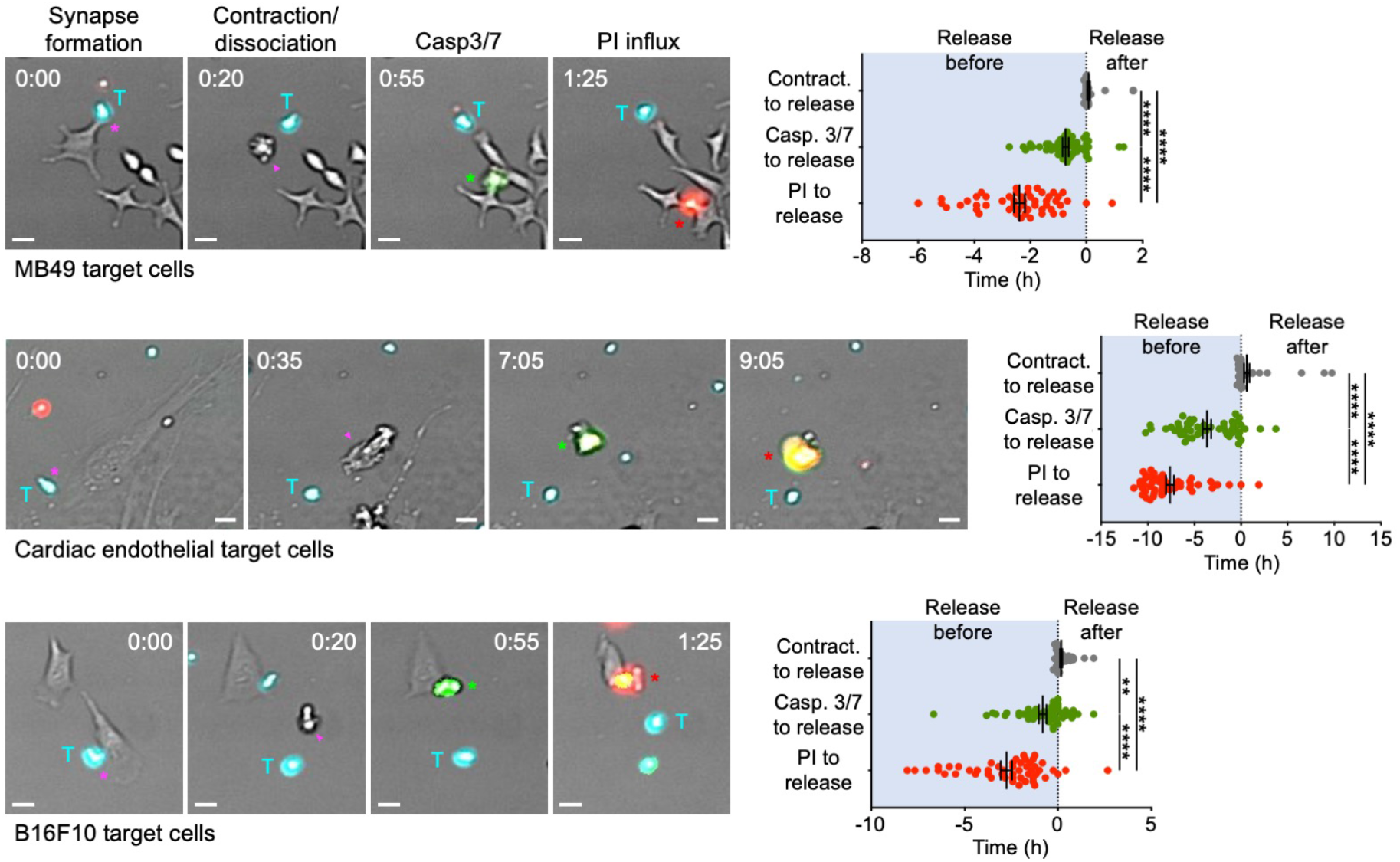
Apoptotic contraction predicts CTL dissociation from multiple target cell types. CTV-labeled OT-1 CTLs were imaged together with the indicated OVA-loaded target cells in the presence of both CellEvent Caspase 3/7 and PI. Left, time-lapse montages of representative contacts. Conjugate formation is indicated by magenta asterisks, dissociation by magenta arrowheads, CellEvent Caspase 3/7 flux by green asterisks, and PI influx by red asterisks. Time in H:MM is shown in the top left or right corner of each image. Scale bars = 20 μm. Right, quantification of offset time between contraction (Contract.), CellEvent Caspase 3/7 flux (Casp. 3/7), or PI flux and target cell release. Data points to the left of the origin denote detachment before the index of interest, whereas data points to the right of the origin denote detachment after. N = 52 MB49 conjugates, 51 endothelial cell conjugates, and 51 B16F10 conjugates. Error bars denote SEM. P-values were calculated by one-way ANOVA with Tukey correction. ** and **** denote P ≤ 0.01 and P ≤ 0.0001, respectively.

**Supplementary figure 3.**
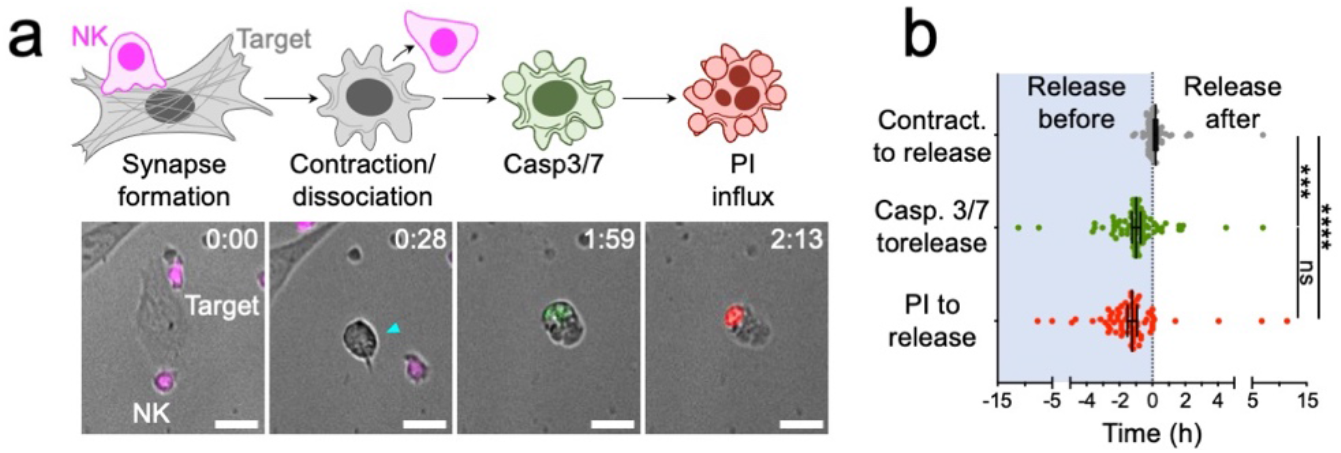
Apoptotic contraction correlates with NK cell dissociation. CTV-labelled, primary murine NK cells were imaged together with B16F10 target cells in the presence of both CellEvent Caspase 3/7 and PI. (**a**) A time-lapse montage of a representative contact is shown below, with schematic diagrams above. Dissociation is indicated by the cyan arrowhead. Time in H:MM is shown in the top right corner of each image. Scale bars = 20 μm. (**b**) Quantification of offset time between contraction (Contract.), CellEvent Caspase 3/7 flux (Casp. 3/7), or PI flux and target cell release. Data points to the left of the origin denote detachment before the index of interest, whereas data points to the right of the origin denote detachment after. N = 71. Error bars indicate SEM. P-values were calculated by one-way ANOVA with Tukey correction. ***, ****, and ns denote P ≤ 0.01, P ≤ 0.0001, and P > 0.05, respectively.

**Supplementary figure 4.**
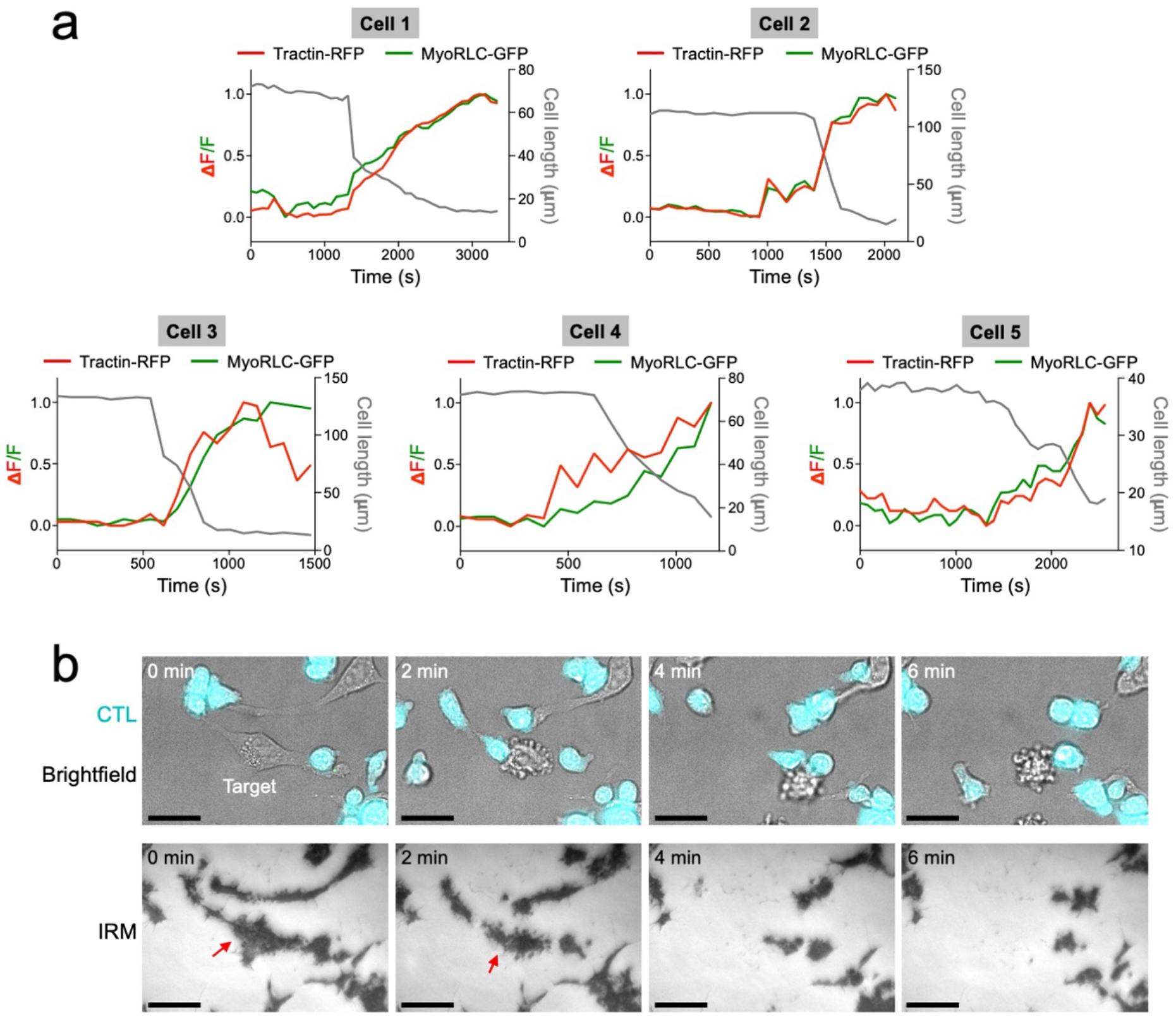
Structural characterization of apoptotic contraction. (**a**) CTV-labeled OT-1 CTLs were imaged together with OVA-loaded MB49 cells expressing both Myl12b-GFP and Tractin-RFP. Mean fluorescence intensity of Myl12b-GFP and Tractin-RFP in five representative target cells is shown, graphed over time together with the length of the cell along its major axis. (**b**) CTV-labeled OT-1 CTLs were imaged with OVA-loaded MB49 cells. A time-lapse montage of a representative killing event is shown, with brightfield and fluorescence channels above and IRM below. Red arrows denote the footprint of the contracting target, which detaches completely from the substrate. Scale bars = 20 μm.

**Supplementary figure 5.**
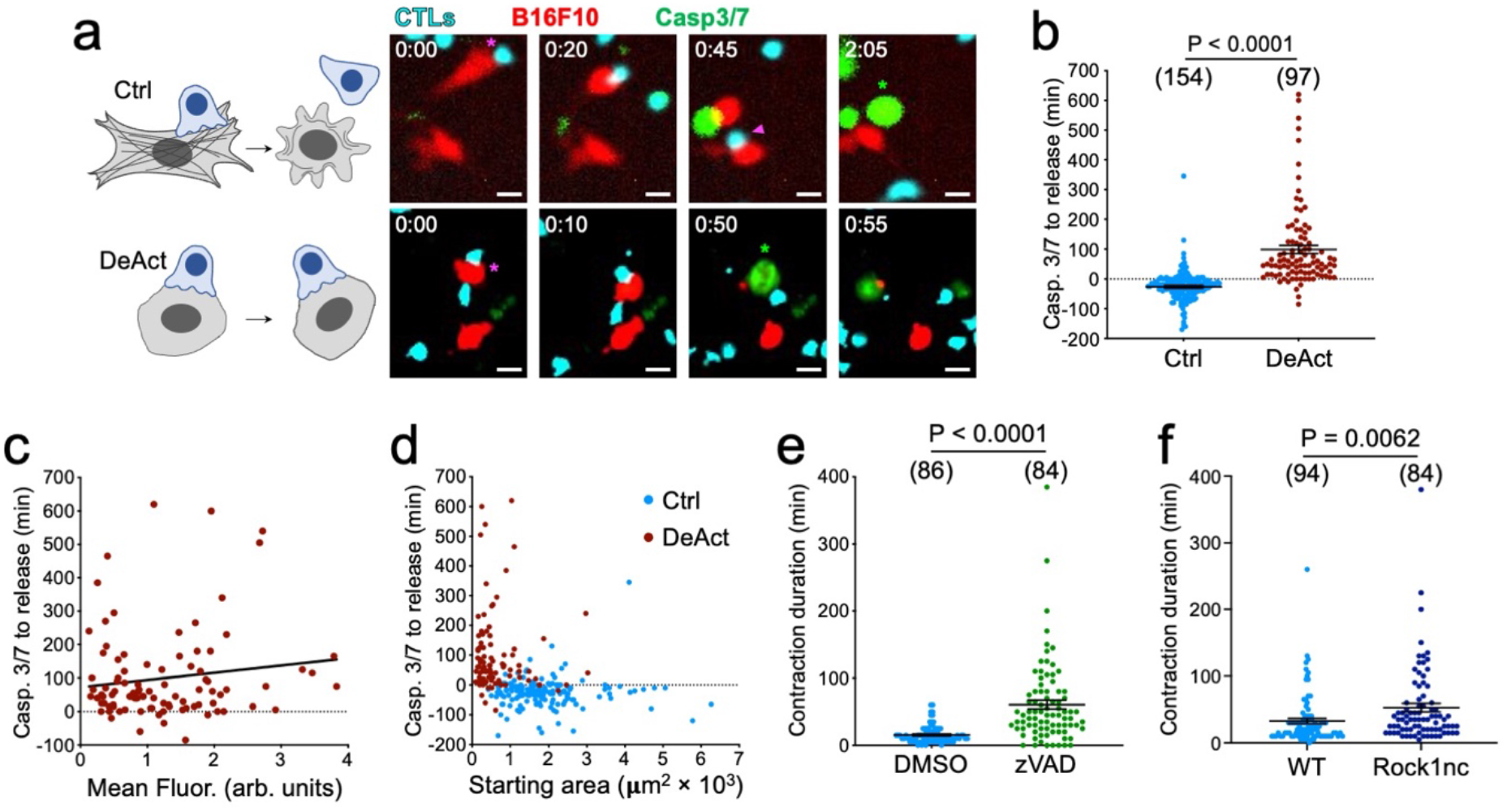
Cytoskeletal contraction is necessary for CTL dissociation. (**a-d**) B16F10 cells expressing Cherry alone (Ctrl) or DeAct together with Cherry (DeAct) were loaded with OVA and imaged with CTV-labelled CTLs in the presence of CellEvent Caspase 3/7. (**a**) Time-lapse montages of representative contacts are shown to the right, with schematic diagrams to the left. Conjugate formation is indicated by magenta asterisks, dissociation by magenta arrowheads, and CellEvent Caspase 3/7 flux by green asterisks. Time in H:MM is shown in the top left corner of each image. Scale bars = 20 μm. (**b**) Quantification of the time delay between CellEvent Caspase 3/7 flux and target cell release. (**c**) 2D plot relating the mean Cherry fluorescence of DeAct expressing targets to the time interval between CellEvent Caspase 3/7 flux and target cell release. (**d**) 2D plot relating the starting area of the indicated target cells to the time interval between CellEvent Caspase 3/7 flux and target cell release. (**e**) The time interval between the start and end of contraction, measured in B16F10 cells engaged by OT-1 CTLs in the presence of 100 μm zVAD-FMK (zVAD) or vehicle control (DMSO). (**f**) The time interval between the start and end of contraction, measured for wild type (WT) and Rock1nc MEFs engaged by OT-1 CTLs. Sample size is shown in parentheses at the top of each column in **b**, **e**, and **f**, with error bars denoting SEM. P-values were calculated by two-tailed unpaired Student’s t test.

**Supplementary figure 6.**
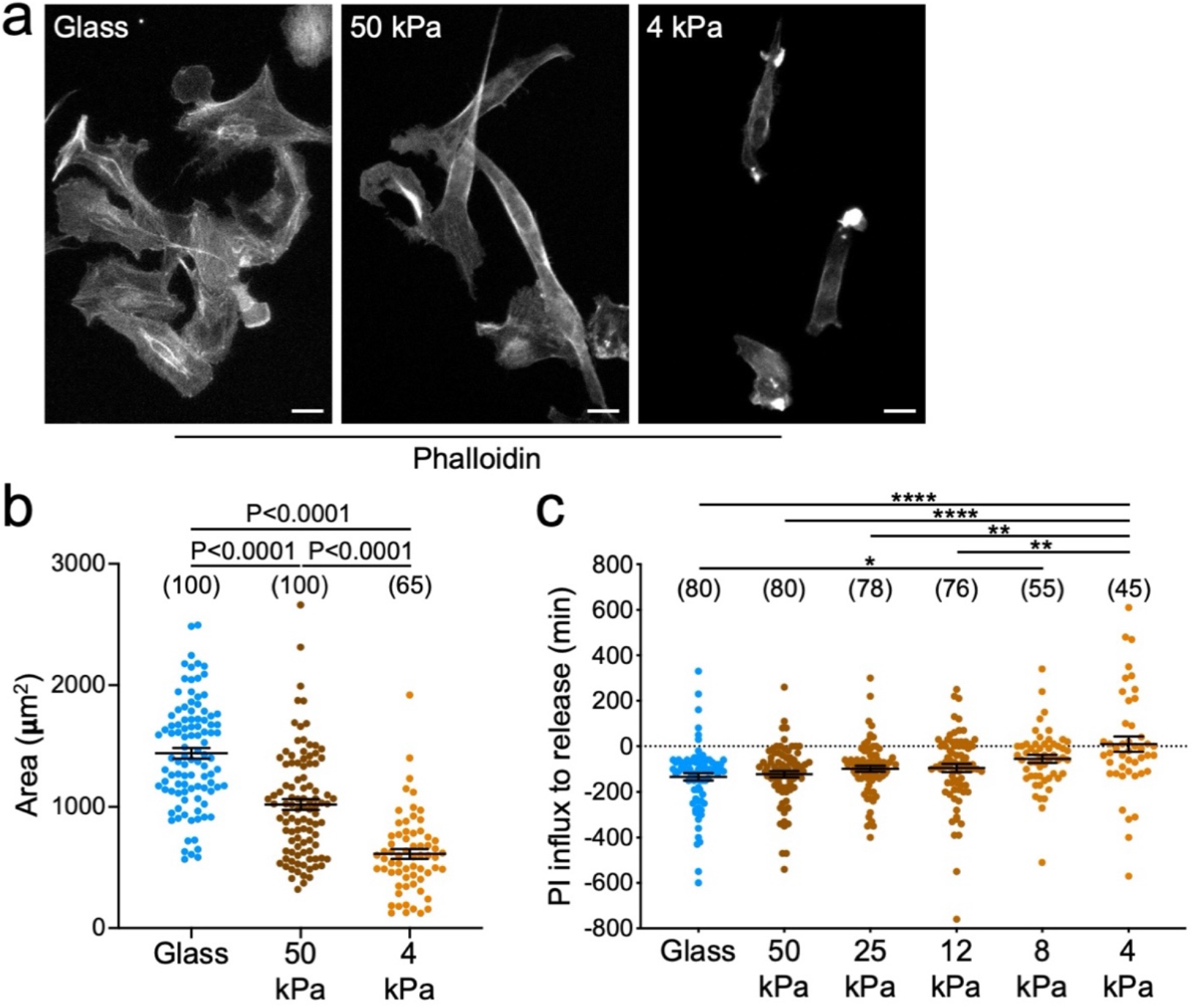
Substrate stiffness controls target cell morphology and CTL dissociation. B16F10 cells were cultured on hydrogels of varying stiffness (4-50 kPa) or glass. (**a**) Representative images of phalloidin stained cells cultured on the indicated surfaces. Scale bars = 20 μm. (**b**) Quantification of target cell area on the indicated substrates. (**c**) Quantification of the time delay between PI influx and target cell release. Sample size is shown in parentheses at the top of each column in **b** and **c**, with error bars denoting SEM. P-values were calculated by one-way ANOVA with Tukey correction. *, **, and **** denote P ≤ 0.05, P < 0.01, and P < 0.0001, respectively.

**Supplementary figure 7.**
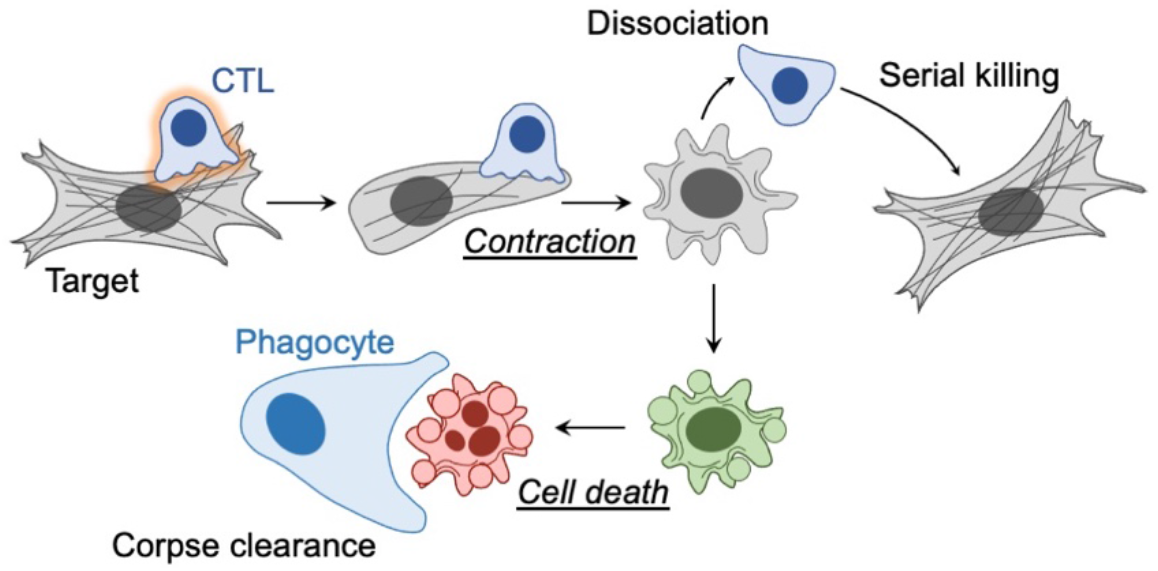
Model for CTL dissociation. Apoptotic contraction promotes CTL detachment by attenuating activating signals. Timely target cell release facilitates both serial killing by the CTL and corpse clearance by patrolling phagocytes.

## Supplementary Movie Legends

**Supplementary movie 1. *Prf1^+/+^* CTLs kill and dissociate from target cells.** CTV-labeled *Prf1^+/+^* (wild type) OT-1 CTLs (cyan) were imaged together with OVA-loaded YFP^+^ B16F10 cells (yellow) in the presence of PI (red). A representative 600× time-lapse movie is shown. Time in HH:MM is shown in the upper left corner.

**Supplementary movie 2. *Prf1^-/-^* CTLs neither kill nor dissociate from target cells.** CTV-labeled *Prf1^-/-^* OT-1 CTLs (cyan) were imaged together with OVA-loaded YFP^+^ B16F10 cells (yellow) in the presence of PI. A representative 600× time-lapse movie is shown. Time in HH:MM is shown in the upper left corner.

**Supplementary movie 3. CTL dissociation is tightly correlated with apoptotic contraction.** CTV-labeled OT-1 CTLs (cyan) were imaged together with OVA-loaded B16F10 cells in the presence of both CellEvent Caspase 3/7 Green (green) and PI (red). A representative 600× timelapse movie is shown. Fluorescence signals have been overlaid onto the brightfield images. Time in HH:MM is shown in the upper right corner.

**Supplementary movie 4. CTL dissociation correlates with apoptotic contraction in 3D culture.** CTV-labeled OT-1 CTLs (cyan) and OVA-loaded YFP^+^ B16F10 cells (yellow) were embedded in collagen and then imaged in the presence of PI (red). A representative 1500× timelapse movie is shown. Fluorescence signals have been overlaid onto the brightfield images. Time in HH:MM is shown in the upper left corner.

**Supplementary movie 5. Contraction-induced dissociation is not specific to the lytic IS.** CTV-labeled *Prf1^+/+^* (wild type) OT-1 CTLs (cyan) and CellVue Maroon-labeled *Prf1^-/-^* OT-1 CTLs (white) were mixed and then imaged together with OVA-loaded YFP^+^ B16F10 cells (yellow) in the presence of PI (red). A representative 600× time-lapse movie is shown. Time in HH:MM is shown in the lower left corner.

**Supplementary movie 6. Actomyosin cables accumulate in contracting target cells.** MB49 target cells expressing Myl12b-GFP (green) and Tractin-RFP (red) were OVA-loaded and then imaged by confocal microscopy together with CTV-labeled OT-1 CTLs (cyan). A representative 308× time-lapse movie is shown. Time in H:MM:SS.SSS is shown in the lower right corner.

**Supplementary movie 7. CTL dissociation from targets under control conditions.** OVA-loaded B16F10 cells were pretreated with vehicle (DMSO) and then imaged together with CTV-labeled OT-1 CTLs (cyan) in the presence of both CellEvent Caspase 3/7 Green (green) and PI (red). A representative 600× time-lapse movie is shown. Fluorescence signals have been overlaid onto the brightfield images. Time in HH:MM is shown in the upper left corner. This control response is included for comparison with the MycB condition (supplementary movie 8).

**Supplementary movie 8. Depletion of target cell F-actin inhibits CTL dissociation.** OVA-loaded B16F10 cells were pretreated with MycB and then imaged together with CTV-labeled OT-1 CTLs (cyan) in the presence of both CellEvent Caspase 3/7 Green (green) and PI (red). A representative 600× time-lapse movie is shown. Fluorescence signals have been overlaid onto the brightfield images. Time in HH:MM is shown in the upper left corner.

**Supplementary movie 9. CTL dissociation from targets under control conditions.** B16F10 cells expressing control Cherry vector (red) were OVA-loaded and then imaged together with CTV-labeled OT-1 CTLs (cyan) in the presence of CellEvent Caspase 3/7 Green (green). A representative 600× time-lapse movie is shown. Time in HH:MM is shown in the upper left corner. This control response is included for comparison with the DeAct condition (supplementary movie 10).

**Supplementary movie 10. Depletion of target cell F-actin inhibits CTL dissociation.** B16F10 cells expressing DeAct and Cherry (red) were OVA-loaded and then imaged together with CTV-labeled OT-1 CTLs (cyan) in the presence of CellEvent Caspase 3/7 Green (green). A representative 600× time-lapse movie is shown. Time in HH:MM is shown in the upper left corner.

**Supplementary movie 11. CTL dissociation from targets under control conditions.** OVA-loaded YFP^+^ B16F10 cells (yellow) were pretreated with vehicle (DMSO) and then imaged together with CTV-labeled OT-1 CTLs (cyan) in the presence PI (red). A representative 960× time-lapse movie is shown. Time in HH:MM is shown in the upper left corner. This control response is included for comparison with the zVAD condition (supplementary movie 12).

**Supplementary movie 12. Caspase inhibition delays CTL dissociation.** OVA-loaded YFP^+^ B16F10 cells (yellow) were pretreated with zVAD and then imaged together with CTV-labeled OT-1 CTLs (cyan) in the presence PI (red). A representative 960× time-lapse movie is shown. Time in HH:MM is shown in the upper left corner.

**Supplementary movie 13. CTL dissociation from targets under control conditions.** OVA-loaded MEFs were imaged together with CTV-labeled OT-1 CTLs (cyan) in the presence of PI (red). A representative 1200× time-lapse movie is shown. Fluorescence signals have been overlaid onto the brightfield images. Time in HH:MM is shown in the upper left corner. This control response is included for comparison with the Rock1nc condition (supplementary movie 14).

**Supplementary movie 14. Rock1 cleavage promotes target cell contraction and CTL dissociation.** OVA-loaded Rock1nc MEFs were imaged together with CTV-labeled OT-1 CTLs (cyan) in the presence of PI (red). A representative 1200× time-lapse movie is shown. Fluorescence signals have been overlaid onto the brightfield images. Time in HH:MM is shown in the upper left corner.

**Supplementary movie 15. Cytoskeletal contraction is sufficient to induce CTL dissociation.** MB49 cells expressing GFP-Rock2-ER (green) were OVA-loaded and then imaged together with CTV-labeled *Prf1^-/-^* OT-1 CTLs (cyan). A representative 600× time-lapse movie is shown. Fluorescence signals have been overlaid onto the brightfield images. Time in HH:MM is shown in the upper right corner. 4-OHT was added to the cells at the 35 min time point to induce GFP-Rock2-ER dimerization and activation.

**Supplementary movie 16. GFP-ER induces neither target contraction nor CTL dissociation.** MB49 cells expressing GFP-ER (green) were OVA-loaded and then imaged together with CTV-labeled *Prf1^-/-^* OT-1 CTLs (cyan). A representative 600× time-lapse movie is shown. Fluorescence signals have been overlaid onto the brightfield images. Time in HH:MM is shown in the lower left corner. 4-OHT was added to the cells at the 40 min time point to induce GFP-ER dimerization.

**Supplementary movie 16. Reduced CTL Ca^2+^ signaling precedes dissociation.** OT-1 CTLs were loaded with Fura-2AM and then imaged together with OVA-loaded B16F10 cells. A representative 600× time-lapse movie is shown. Fura-2 ratio is depicted in pseudocolor with cold and warm colors indicating low and high intracellular Ca^2+^, respectively. Fura-2 signals have been overlaid onto the brightfield images. Time in HH:MM is shown in the upper left corner.

